# Psilocybin facilitates fear extinction: importance of dose, context, and serotonin receptors

**DOI:** 10.1101/2024.05.04.592469

**Authors:** Samuel C. Woodburn, Caleb M. Levitt, Allison M. Koester, Alex C. Kwan

**Author notes:** Correspondence to Alex Kwan, Ph.D., Room 111 Weill Hall, 526 Campus Road, Ithaca, NY, 14853, United States.

## Abstract

A variety of classic psychedelics and MDMA have been shown to enhance fear extinction in rodent models. This has translational significance because a standard treatment for posttraumatic stress disorder (PTSD) is prolonged exposure therapy. However, few studies have investigated psilocybin’s potential effect in fear learning paradigms. More specifically, the extents to which dose, timing of administration, and serotonin receptors may influence psilocybin’s effect on fear extinction are not understood. In this study, we used an auditory delay fear conditioning paradigm to determine the effects of psilocybin on fear extinction, extinction retention, and fear renewal in male and female mice. Psilocybin robustly enhances fear extinction when given acutely prior to testing for all doses tested. Psilocybin exerts long-term effects to elevate extinction retention and suppress fear renewal in a novel context, though these changes were sensitive to dose. Administration of psilocybin prior to fear learning or immediately after extinction yielded no change in behavior, indicating that concurrent extinction experience is necessary for the drug’s effects. Co-treatment with a 5-HT_2A_ receptor antagonist blocked psilocybin’s effects for extinction, extinction retention and fear renewal, whereas 5-HT_1A_ receptor antagonism attenuated only the effect on fear renewal. Collectively, these results highlight dose, context, and serotonin receptors as crucial factors in psilocybin’s ability to facilitate fear extinction. The study provides preclinical evidence to support investigating psilocybin as a pharmacological adjunct for extinction-based therapy for PTSD.

## INTRODUCTION

Post-traumatic stress disorder (PTSD) is a debilitating condition in which a traumatic experience causes difficulty in distinguishing safe and unsafe contexts, resulting in exaggerated responses to stimuli reminiscent of the initial trauma^1^. A standard treatment for PTSD is prolonged exposure therapy, wherein patients learn to extinguish fearful responses through repeated presentations of triggering stimuli in a safe setting^2^. While generally effective, many individuals discontinue the therapy due to the challenging emotional reactions upon re-exposure^3^. Further, extinction learning is often bound to the context in which it was learned^4–6^. In other words, patients may respond well in the clinic but continue to experience PTSD symptoms in their daily lives^2^. To overcome these limitations, a considerable amount of research has focused on developing pharmacotherapies that can promote sustained and context-independent extinction^7– 9^.

Classic psychedelics and related psychoactive compounds may have therapeutic potential for mental illnesses including PTSD^10–12^. The promise is highlighted by the Phase 3 clinical trials of 3,4-methylenedioxymethamphetamine (MDMA), where patients reported an enduring decrease in PTSD symptoms following MDMA-assisted psychotherapy^13,14^. Consistently, in preclinical mouse models involving auditory cued fear conditioning, a single dose of MDMA reduced conditioned freezing 30 minutes after administration^15^. Importantly, this behavioral change lasted 10 days later and persisted when mice were tested in an unfamiliar context. The enhancing effect of MDMA on extinction was moderated by dose^15^ and required 5-HT_2A_ receptors^16^. Recently, classic psychedelics have been evaluated using preclinical mouse models for fear learning. For instance, mice treated with 2,5-dimethoxy-4-iodoamphetamine (DOI) showed less cue-conditioned freezing during fear extinction compared to controls 30 minutes after administration^17^. However, these effects were lost when the mice were tested again 24 hours later, suggesting that the effect of DOI on extinction was not retained. Similar acute, immediate effects on cued fear extinction have been demonstrated for N,N-dimethyltryptamine (DMT) in rats^18,19^, 4-hydroxy-diisopropyltryptamine (4-OH-DiPT) in mice^20^, and TCB-2 in mice^21^. However, the studies so far with classic psychedelics typically have limited scope: testing a single dose, focusing on only acute effects, or omitting fear renewal, making it unclear if the classic psychedelics can have enduring impact that is retained outside the initial extinction context.

Among classic psychedelics, psilocybin has advanced the most in clinical trials. Recent work indicates that psilocybin administration with psychological support may be an effective treatment option for major depression and treatment-resistant depression^22–25^. Excitingly, some clinical reports show reductions in symptom severity months after treatment^26,27^, suggesting that psilocybin-based interventions promote long-term behavioral changes. It has been suggested that psilocybin may have therapeutic potential for PTSD as well^28,29^. In a small open-label study, PTSD severity declined in long-term AIDS survivors following psilocybin-assisted group psychotherapy^30^. This early result is now followed by ongoing clinical trials of psilocybin in people with trauma-related disorders from frontline clinical duty (NCT05163496), experience in adulthood (NCT05312151), and combat for veterans (NCT05554094).

Less is known about the effects of psilocybin in rodent models of fear learning. In one report, mice given low dose (0.1 - 0.5 mg/kg) psilocybin prior to fear conditioning extinguished conditioned freezing more quickly than controls^31^. However, this dosing strategy makes it difficult to discern whether psilocybin’s effect was due to a change in extinction learning or in the consolidation and retrieval of the initial fear memory. More recent studies suggest that psilocybin (1 - 2 mg/kg) reduces conditioned freezing when administered 30 minutes prior to extinction^32,33^, and that this effect can persist 24 hours later^32^, but used contextual and trace fear conditioning paradigms. Thus, despite tantalizing hints from these recent studies, much remains unknown regarding psilocybin’s effect on cued fear extinction. Crucially, the extent to which psilocybin’s effects may depend on essential parameters such as dose, timing of administration, and serotonin receptors are not well understood.

In this study, we used auditory delay fear conditioning and determined the effects of psilocybin on fear extinction, extinction retention, and fear renewal with a total of 112 male and female mice. We report that a single 1 mg/kg dose of psilocybin enhances fear extinction and suppresses fear renewal in a novel context up to 8 days after drug administration. We found that the persisting action of psilocybin is sensitive to dose, despite its acute impact being observed at all doses tested. We varied the timing of administration to show that psilocybin was ineffective at altering the acquisition of conditioned fear or consolidation of fear extinction. We tested the role of 5-HT_1A_ and 5-HT_2A_ receptors in supporting psilocybin’s effects on fear extinction and renewal. Collectively, these results demonstrate that psilocybin facilities fear extinction in mice and delineate the parameters under which its behavioral effects occur.

## MATERIALS AND METHODS

### Animals

Male and female C57BL/6 mice were purchased from Jackson Laboratory (Stock No. 000664) and given 14 days to acclimate to the vivarium at Cornell University. Within the vivarium, animals were kept in a climate-controlled room and were housed in groups of 4 mice per cage with *ad libitum* access to food and water. Mice were kept on a 12-hour light/dark cycle (lights on between 08:00 and 16:00 hours), with behavioral testing occurring between 08:30 to 13:00 hours. All mice were randomly assigned to different experimental groups prior to testing. Animals were 7 - 8 weeks old at the start of behavioral studies. Animal care and experimental procedures were approved by the Institutional Animal Care & Use Committee (IACUC) at Cornell University.

### Drugs

Psilocybin was obtained from the Usona Institute’s Investigational Drug & Material Supply Program. The chemical composition of psilocybin was confirmed by high performance liquid chromatography at Usona Institute. A stock 4 mg/mL solution was prepared by dissolving solid in 0.9% saline, then diluted into working solutions of 0.05, 0.1, or 0.2 mg/mL the day before injection. The 5-HT_1A_ receptor antagonist WAY100635 (4380, Tocris Biosciences) was diluted in 0.9% saline for a 5 mg/mL stock solution. The 5-HT_2A_ receptor antagonist MDL100907 (4173, Tocris Biosciences) was dissolved in a 0.9% saline solution containing 0.3% Tween 80 (P1754, Sigma-Aldrich) for a 1 mg/mL stock solution. The day before injection, aliquots of WAY100635 or MDL100907 were diluted in 0.9% saline to produce working solutions of 0.2 mg/mL and 0.1 mg/mL respectively. For injections of two drugs, cocktails of psilocybin (0.1 mg/mL) and either WAY100635 (0.2 mg/mL) or MDL100907 (0.1 mg/mL) were prepared using the same stock solutions and diluting with 0.9% saline. This was done so that both drugs could be administered as a single injection, mitigating stress induced by performing multiple injections. Injections, including vehicle controls, were performed intraperitoneally (i.p.) at a volume of 10 mL/kg. Methods for drug preparation and doses of psilocybin (0.5, 1, or 2 mg/kg), WAY100635 (2 mg/kg), and MDL100907 (1 mg/kg) were chosen based on prior work^33–35^.

### Fear conditioning and extinction paradigms

The timeline for delay fear conditioning and extinction tests was inspired by prior literature^15,17,18^. All behavioral testing was conducted in a near-infrared video fear conditioning system (MED-VFC2-SCT-M, Med Associates Inc). Prior to each session, mice were brought into a side room separated from the fear conditioning system where they habituated for 30 minutes. The near-infrared video camera was calibrated to record with the manufacturer’s recommended average intensity of 130 a.u. so that conditions for motion detection were held constant across all experimental days. On day 1 of testing (auditory cued fear conditioning), the mouse was individually placed in the conditioning chamber, which was prepared in “context A” (blank straight walls, metal grid floor, cleaned with 70% ethanol before each session). Each conditioning trial began with a 3-minute habituation period, after which the mouse was given 5 presentations of an auditory tone as the conditioned stimulus (CS; 4 kHz, 80 dB, 30 s duration). Each CS co-terminated with a foot shock unconditioned stimulus (US; 0.8 mA, 2 s duration). A 90 s intertrial interval separated the CS + US pairings. On day 2, mice were left undisturbed in their home cages. On day 3 (fear extinction), the mouse received an i.p. injection of vehicle or drug solution and then were returned to their home cages for 30 minutes. After this, the mouse was placed in the conditioning chamber prepared in “context B” (black A-frame, white plastic floor, cleaned with 1% acetic acid before each session). A 3-minute habituation period was given, followed by 15 presentations of the CS without a US, each of which was separated by a 15 s intertrial interval. On day 4 (extinction retention), the mouse underwent the same procedure as day 3, except with no injection administered beforehand.

Mice were then left undisturbed until day 11 (fear renewal), at which point the mouse was placed in the conditioning chamber prepared in “context C” (striped curved white wall, striped plastic floor, cleaned with Peroxigard™ before each session) and subjected to the same habituation and CS-only schedule used on days 3 and 4. On each experimental day, mice were returned to their housing location immediately after testing. Variations of this general paradigm were performed to determine the behavioral mechanisms supporting psilocybin’s effects. In one set of these experiments (**Fig. 3A**), the mouse was administered 0.9% saline (10 mL/kg, i.p.) or psilocybin (1 mg/kg, i.p.) then returned to their home cage for 30 minutes, after which they underwent fear conditioning followed by extinction 48 hours later as described above. In another set of experiments (**Fig. 3D**), the mouse underwent fear conditioning on day 1, then fear extinction on day 3 with 0.9% saline (10 mL/kg, i.p.) or psilocybin (1 mg/kg, i.p.) administered immediately after fear extinction, then tested for extinction retention 24 hours later.

### Analysis

Conditioned freezing was used as the primary readout for behavior, and was defined as a lack of detectable movement other than breathing for ≥ 1 s. The videos captured by the near-infrared camera were analyzed with automated procedures using the VideoFreeze™ software from Med Associates Inc (motion threshold: 18 au, detection method: linear, minimum freeze duration: 30 frames = 1 s). For extinction, extinction retention, and fear renewal, we binned the data to calculate the freezing responses, where bin “Hab” corresponds to the 3-minute habituation period with no tone, bin 1 corresponds to CS period during tones 1-3, bin 2 corresponds to CS period during tones 4-6, bin 3 corresponds to CS period during tones 7-9, bin 4 corresponds to CS period during tones 10-12, and bin 5 corresponds to CS period during tones 13-15.

### Statistics

Statistical testing was conducted in SPSS. Conditioned freezing data were analyzed with generalized linear mixed effects modeling. Fixed factors are listed in the corresponding figure legends. Random factors of subject, cage, and sex were used to account for individual differences in the overall model, and a heterogeneous autoregressive 1 (ARH1) covariance structure was used in accounting for repeated measures. A sequential Bonferroni correction was applied to between-groups comparisons.

## RESULTS

### Psilocybin enhances extinction and attenuates renewal of conditioned fear

Motivated by studies reporting the effects of MDMA on fear extinction^15,16^, we implemented a delay fear conditioning protocol to test the effects of psilocybin on fear extinction (**Fig. 1A**; see **Materials and Methods**). Briefly, on day 1, we trained a C57BL/6J mouse to associate an auditory tone (conditioned stimulus; CS) with a noxious foot shock (unconditioned stimulus; US) in context A. On day 3, the mouse was given psilocybin (0.5, 1, or 2 mg/kg, i.p.) or saline vehicle. The animal was then placed back in its home cage for 30 minutes before being moved to context B for fear extinction, which consisted of 15 CS presentations without the US pairing. On day 4, retention of the extinction memory was tested by exposing the mouse to another 15 CS presentations in context B. Finally, on day 11, the mouse was exposed to another 15 CS presentations in context C to determine if its conditioned fear would be renewed in an unfamiliar environment distinct from the initial extinction context. For each drug condition, we tested 4 male and 4 female animals. The results are shown in **Figs. 1B-J**.

**Figure 1.**
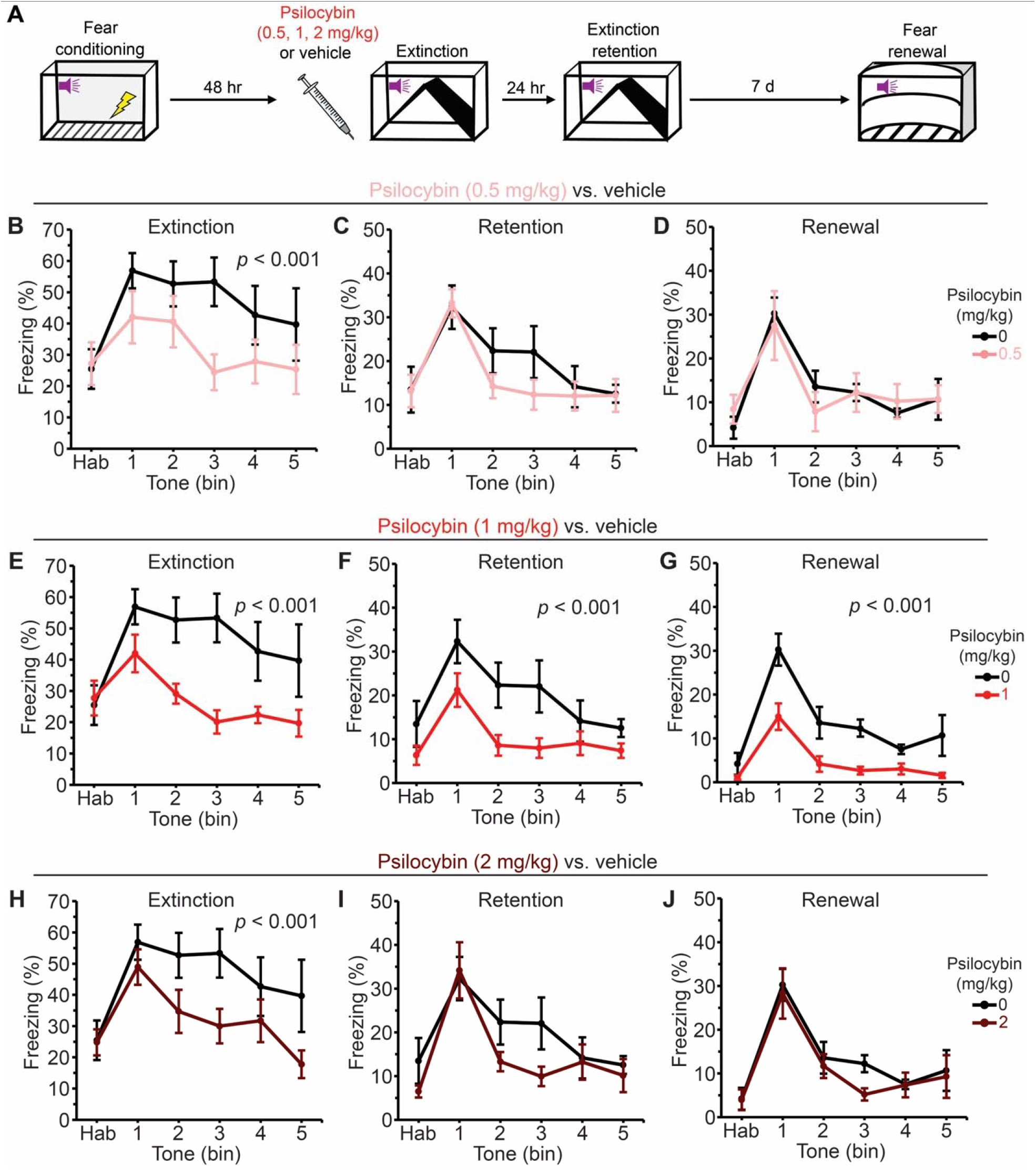
Psilocybin drives persisting enhancement of fear extinction and suppression of fear renewal. **(A)** Experimental timeline. **(B)** Fraction of time spent freezing during habituation (Hab) and different binned CS periods during extinction on day 3. Black, vehicle. Red, 0.5 mg/kg psilocybin. Line, mean. Error bars, ± s.e.m. Shown is the p-value (if <0.05) for between-group post hoc analysis between vehicle and 0.5 mg/kg psilocybin performed after generalized linear mixed effects model with fixed factors of dose, time, and 2-way interactions. **(C)** Similar to (B) for extinction retention on day 4. **(D)** Similar to (B) for fear renewal on day 11. **(E – G)** Similar to (B) – (D) for 1 mg/kg psilocybin. The data for vehicle is repeated from (B) – (D) for purpose of comparison. **(H – J)** Similar to (B) – (D) for 2 mg/kg psilocybin. The data for vehicle is repeated from (B) – (D) for purpose of comparison. N = 32, including 4 males and 4 females for vehicle, and 4 males and 4 females for each of the psilocybin conditions.

For day 3, as expected for an extinction session, freezing response decreased as tones were repeatedly presented (main effect of time: F_12, 420_ = 2.568, *p* = 0.001; generalized linear mixed effects model with fixed factors of dose, time, and 2-way interactions). Psilocybin reduced conditioned freezing (main effect of dose: F_3, 420_ = 6.223, *p* < 0.001), which was statistically significant for all doses tested compared to vehicle controls (*p* < 0.001 in all cases; between-group post hoc analyses), with no psilocybin dose significantly outperforming the others (**Figs. 1B, E, H**). By contrast, for retention on day 4, we detected the effect of psilocybin (main effect of dose: F_3, 420_ = 4.387, *p* = 0.005; main effect of time: F_12, 420_ = 4.998, *p* < 0.001), however post hoc analyses revealed only mice previously given 1 mg/kg psilocybin froze less than vehicle controls (*p* < 0.001), while no significant differences were found between mice given 0.5 or 2 mg/kg psilocybin and vehicle (**Figs. 1C, F, I**). For fear renewal on day 11, we observed a drug effect again (main effect of dose: F_3, 420_ = 5.72, *p* < 0.001; main effect of time: F_12, 420_ = 5.665; *p* < 0.001), which was driven by mice in the 1 mg/kg psilocybin group, as these mice froze less than their counterparts injected with vehicle (*p* = 0.001), 0.5 (*p* = 0.005), or 2 mg/kg psilocybin (*p* = 0.022) (**Figs. 1D, G, J**). Together, the data indicate that psilocybin robustly enhances fear extinction when given acutely prior to testing. Psilocybin also exerts long-term effects to elevate extinction retention and suppress fear renewal up to 8 days later, though these changes were sensitive to dose and were most reliably elicited by 1 mg/kg of psilocybin.

To explore potential difference in male versus female animals, we analyzed the dose-response data using fixed factors of sex, dose, time, and all 2- and 3-way interactions. For fear acquisition, we found a main effect of time (F_3, 152_ = 193.417; *p* < 0.001), but no sex effects were detected (**Fig. 2A, B**). This changed in the extinction task (**Fig. 2C**), which showed a sex × dose interaction (F_3, 360_ = 3.018; *p* = 0.03) and a main effect of time (F_14, 360_ = 2.965; *p* < 0.001). For extinction retention, we detected main effects of dose (F_3, 360_ = 5.298; *p* = 0.001) and time (F_14, 360_ = 4.915; *p* < 0.001) (**Fig. 2D**). Finally, for fear renewal, there was a significant sex × dose × time interaction (F_42, 360_ = 1.42; *p* = 0.05) (**Fig. 2E**). These additional analyses suggest that sex may be a factor in psilocybin’s dose-dependent effects on fear extinction.

**Figure 2.**
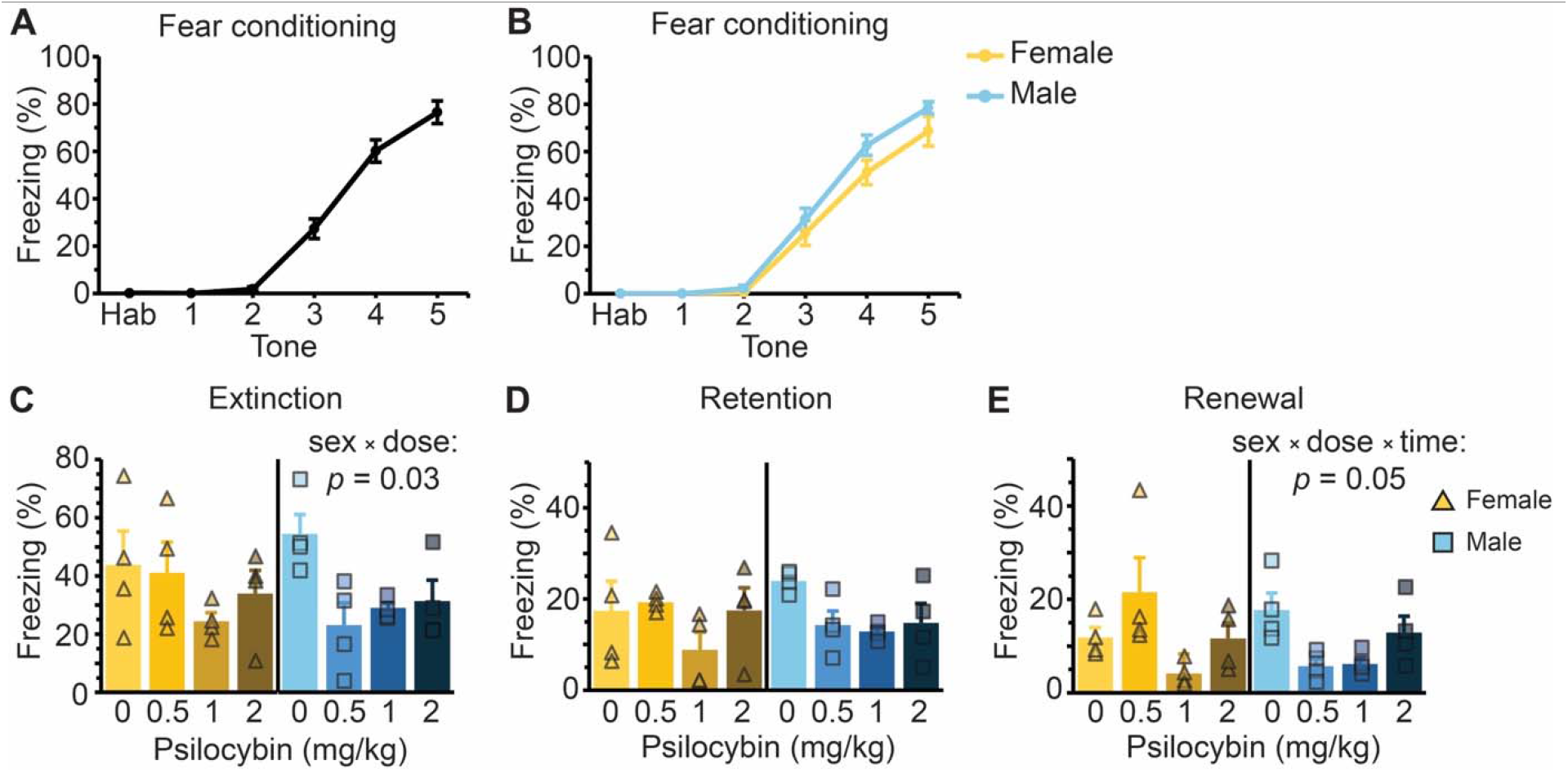
Interaction of dose and sex on psilocybin-enhanced fear extinction. **(A)** Fraction of time spent freezing during different CS presentations during fear conditioning on day 1, for all animals tested in Fig. 1 (N = 32). There was a main effect of time (F_3, 144_ = 172.135; *p* < 0.001). Post hoc analyses did not find any pre-existing difference in freezing between treatment groups prior to injection. Line, mean. Error bars, ± s.e.m. **(B)** Similar to (A) except plotted separately for females (yellow; N = 16) and males (blue; N = 16). No sex difference was detected. **(C)** Fraction of time spent freezing in all CS presentations during extinction on day 3, with females (yellow) and males (blue) plotted separately for each dosing group (n = 4 per sex and dose). Each triangle or square represents an individual animal. Bar, mean. Error bars, ± s.e.m. **(D)** Similar to (C) for extinction retention on day 4. **(E)** Similar to (C) for fear renewal on day 11. For statistical test, we used a generalized linear mixed effects model with fixed factors of sex, dose, time, and all 2- and 3-way interactions.

### Psilocybin is only effective when administered prior to extinction context

How can psilocybin facilitate extinction? Classic psychedelics may boost learning through their effects on neural plasticity, which have been demonstrated for psilocybin as it evokes structural rewiring in the medial frontal cortex^34^ and induces plasticity-related gene expression^37,38^. Given that learning and memory are linked to structural plasticity^39–42^, one possibility here is that learning is enhanced broadly by psilocybin. If true, psilocybin should also improve fear memory acquisition and extinction memory consolidation. An alternative possibility is that learning is enhanced but only for specific contexts. Preclinical studies involving MDMA indicate that the context immediately after drug administration is an important factor in determining the eventual behavioral changes in mice^15^, but it is unknown if this applies to psilocybin and extinction.

To disambiguate the possibilities, we devised two variants of the conditioned fear extinction protocol. For the first variant, mice received 1 mg/kg psilocybin or vehicle 30 minutes prior to fear conditioning on day 1, then underwent fear extinction on day 3 (**Fig. 3A**). As expected, mice readily acquired conditioned freezing during fear conditioning (main effect of time: F_4, 70_ = 105.696, *p* < 0.001; **Fig. 3B**) and progressively reduced conditioned freezing during extinction (main effect of time: F_14, 210_ = 2.641, *p* < 0.001; **Fig. 3C**). However, no difference was observed between the psilocybin and vehicle-treated groups.

**Figure 3.**
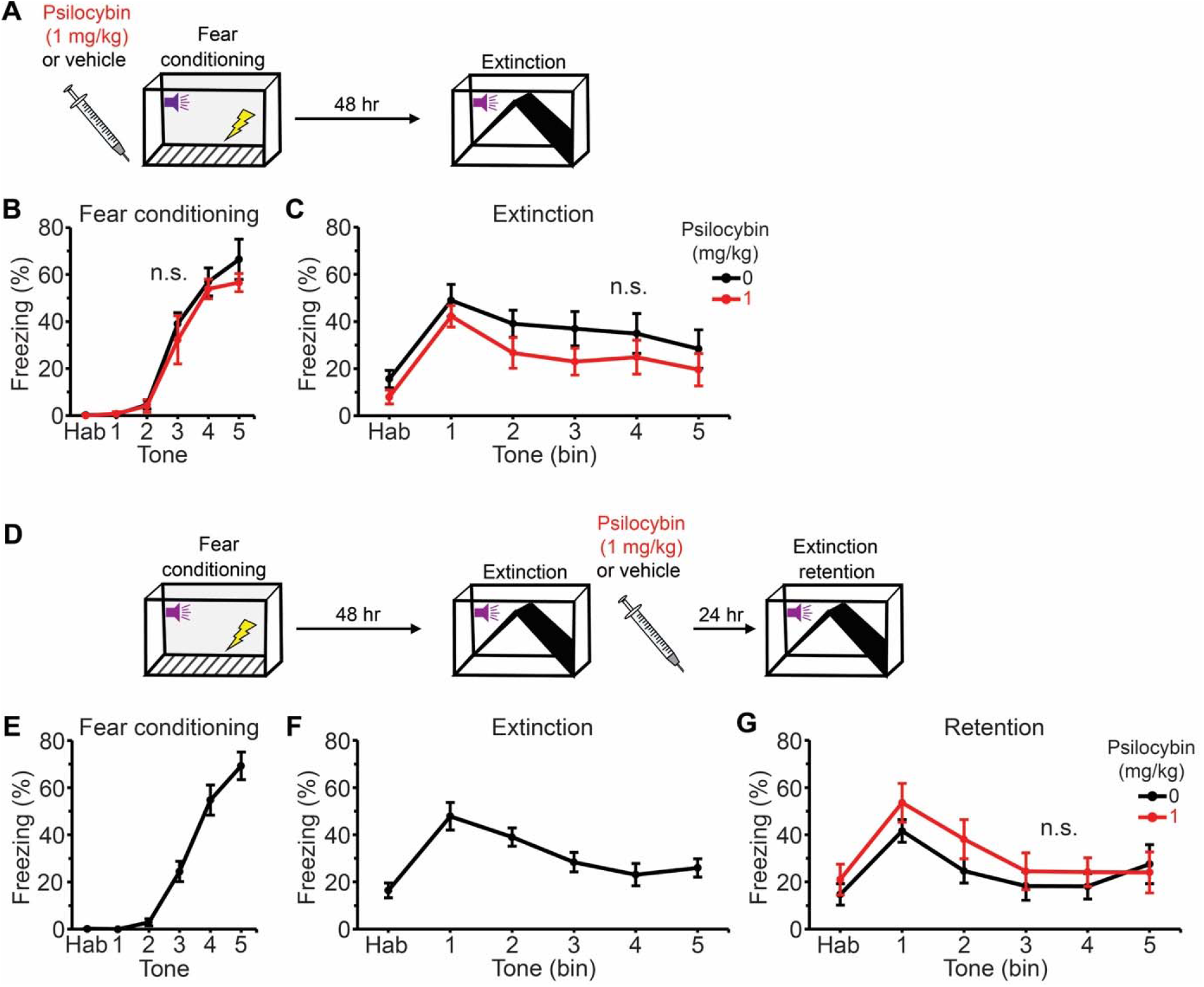
Psilocybin does not enhance fear acquisition nor extinction consolidation. **(A)** Experimental timeline to test psilocybin’s effect on fear acquisition. **(B)** Fraction of time spent freezing during different CS presentations during fear conditioning on day 1. Black, vehicle. Red, 1 mg/kg psilocybin. Line, mean. Error bars, ± s.e.m. n.s., not significant for between-group post hoc analysis between vehicle and psilocybin performed after generalized linear mixed effects model with fixed factors of dose, time, and 2-way interactions. **(C)** Similar to (B) for freezing during habituation and different binned CS periods during extinction on day 3. **(D)** Experimental timeline to test psilocybin’s effect on extinction consolidation. **(E)** Fraction of time spent freezing during different CS presentations during fear conditioning on day 1, for all animals. Line, mean. Error bars, ± s.e.m. **(F)** Similar to (E) for freezing during habituation and different binned CS periods during extinction on day 3. **(G)** Fraction of time spent freezing during different CS presentations during extinction retention on day 4. Black, vehicle. Red, 1 mg/kg psilocybin. Line, mean. Error bars, ± s.e.m. N = 16, including 4 males and 4 females for each of the vehicle and psilocybin conditions for fear acquisition, 4 males and 4 females for each of the vehicle and psilocybin conditions for extinction retention.

For the second variant, mice underwent fear conditioning on day 1 (**Fig. 3D**). Instead of administering psilocybin prior to extinction on day 3, the paradigm was modified such that 1 mg/kg psilocybin or vehicle was administered immediately after extinction. The mice were then tested for extinction retention on day 4. Again, mice readily acquired conditioned freezing during fear conditioning (main effect of time: F_3, 70_ = 63.946; *p* < 0.001; **Fig. 3E**) and progressively reduced conditioned freezing during extinction (main effect of time: F_14, 210_ = 3.275; *p* < 0.001; **Fig. 3F**), with no pre-existing differences between treatment groups prior to drug administration. The post-extinction administration of 1 mg/kg psilocybin did not impact extinction retention on day 4 compared to vehicle controls (**Fig. 3G**). Collectively, these experiments suggest that the timing of psilocybin administration – shortly prior to the initial extinction experience – is key to its effect on retention of the extinction memory.

### 5-HT_2A_ and 5-HT_1A_ receptors contribute to psilocybin’s long-term behavioral effects

Upon entering the body, psilocybin is rapidly converted to its active metabolite, psilocin, which is an agonist at various serotonin receptors^43^, including the 5-HT_1A_ and 5-HT_2A_ subtypes which are highly expressed in the neocortex^44^. To test the role of 5-HT_1A_ and 5-HT_2A_ receptors in mediating psilocybin’s effects on fear extinction, we modified the auditory fear conditioning protocol such that the mouse received an i.p. injection of a mixture containing MDL100907 (1 mg/kg; 5-HT_2A_ receptor antagonist^45,46^), WAY100635 (2 mg/kg; 5-HT_1A_ receptor antagonist^47^) or vehicle, paired with psilocybin (1 mg/kg) or vehicle (**Fig. 4A**). For each of the 4 conditions in MDL100907/vehicle + psilocybin/vehicle, we tested 4 male and 4 female animals. For each of the 4 conditions in WAY100635/vehicle + psilocybin/vehicle, we tested 4 male and 4 female animals. We did not find any notable difference in the results involving the vehicle + psilocybin/vehicle conditions, therefore combined the data to produce **Figs. 4B-D**. The remainder of the results are shown separately for MDL100907 + psilocybin/vehicle (**Figs. 4E-G**) and WAY100635 + psilocybin/vehicle (**Figs. 4H-J**).

**Figure 4.**
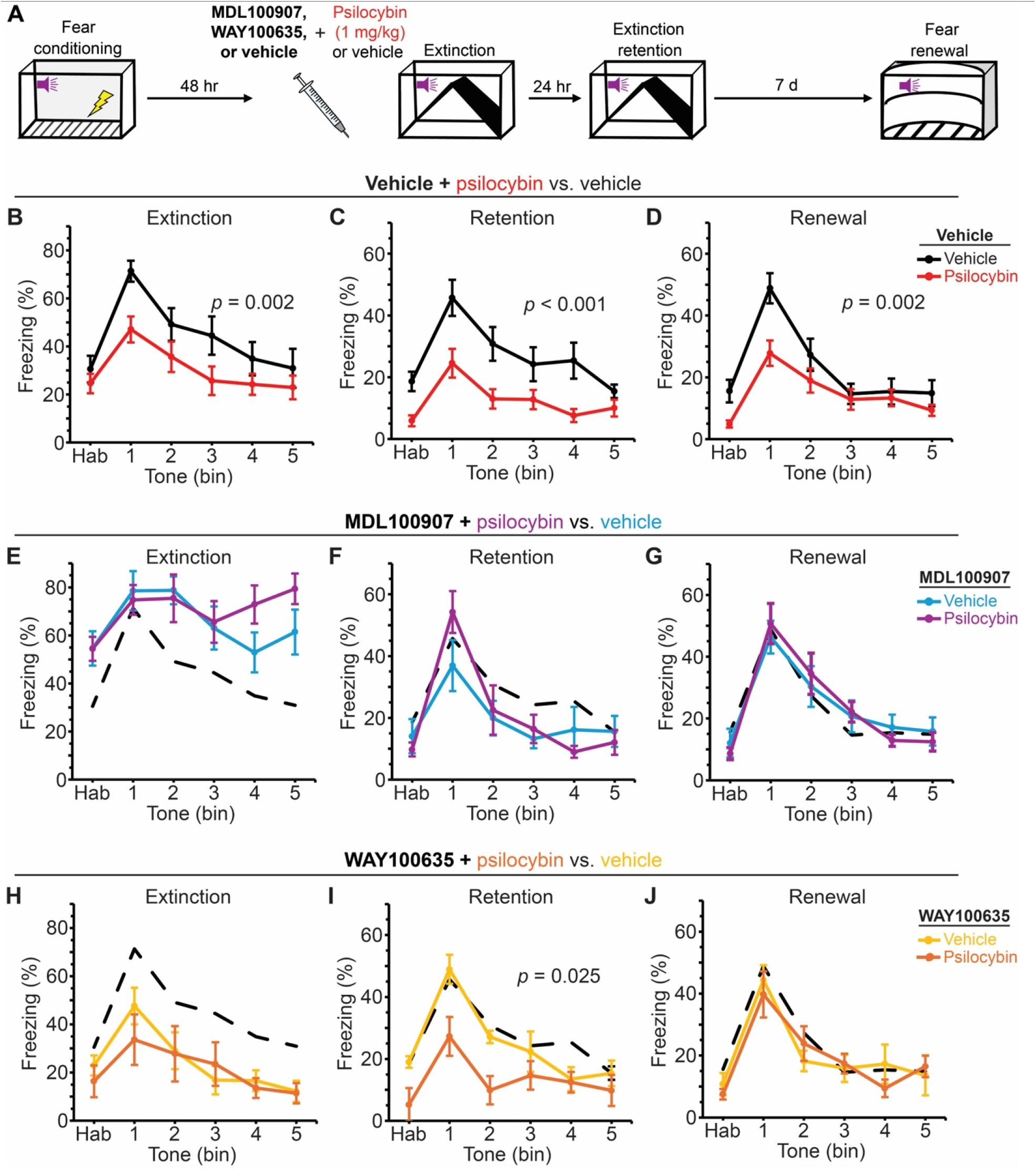
Effects of 5-HT_1A_ and 5-HT_2A_ receptor antagonists on psilocybin-enhanced fear extinction. **(A)** Experimental timeline. **(B)** Fraction of time spent freezing during habituation and different binned CS periods during extinction on day 3. Black, vehicle + vehicle. Red, vehicle + 1 mg/kg psilocybin. Line, mean. Error bars, ± s.e.m.. Shown is the p-value (if <0.05) for between-group post hoc analysis between vehicle + vehicle and vehicle + 1 mg/kg psilocybin performed after generalized linear mixed effects model with fixed factors of antagonist, psilocybin, time, as well as all 2- and 3-way interactions. **(C)** Similar to (B) for extinction retention on day 4. **(D)** Similar to (B) for fear renewal on day 11. **(E – G)** Similar to (B) – (D) for MDL100907 + vehicle (teal) and MDL100907 + psilocybin (purple). Dashed line, the vehicle + vehicle data repeated from (B) – (D) for purpose of comparison. **(H – J)** Similar to (B) – (D) for WAY100635 + vehicle (yellow) and WAY100635 + psilocybin (orange). Dashed line, the vehicle + vehicle data repeated from (B) – (D) for purpose of comparison. N = 64 including 8 males and 8 females for vehicle + vehicle, 8 males and 8 females for vehicle + psilocybin, and 4 males and 4 females each for MDL100907 + vehicle, MDL100907 + psilocybin, WAY100635 + vehicle, and WAY100635 + psilocybin.

For extinction on day 3, the co-administration of a receptor antagonist altered the conditioned freezing response (antagonist × drug interaction: F_2, 870_ = 3.604, *p* = 0.028; generalized linear mixed effects model). Replicating our earlier result in **Fig. 1E**, vehicle + psilocybin mice froze significantly less than vehicle + vehicle controls (*p* = 0.002; **Fig. 4B**). Mice with WAY100635, regardless of whether they received psilocybin, froze less than vehicle + vehicle controls (*p* = 0.001; **Fig. 4E**). Conversely, mice with MDL100907, regardless of whether they received psilocybin, increased freezing compared to vehicle + vehicle controls (*p* = 0.001; **Fig. 4H**). Our interpretation is that, at the doses used, the receptor antagonists themselves significantly altered the measurement of freezing response. Indeed, it has been shown that WAY100635 increases rearing in light box^47^ and MDL100907 reduces spontaneous motor activity in rats and mice^46^, which would affect the readout by our automated video analysis. In agreement with this explanation, MDL100907-treated mice froze more than any other group during the 3-minute habituation period prior to CS presentation (*p* < 0.01 in all cases; bin “Hab” in **Fig. 4E**). Notwithstanding the confound of acute locomotor effects, mice that received receptor antagonists appeared awake on day 3, and thus were expected to have still formed an extinction memory.

For extinction retention on day 4, there was a significant antagonist × psilocybin × time interaction (F_28, 870_ = 1.791; *p* = 0.007). Replicating our previous finding in **Fig. 1F**, vehicle + psilocybin mice froze less than their vehicle + vehicle treated counterparts (*p* < 0.001; **Fig. 4C**). MDL100907 + psilocybin and MDL100907 + vehicle animals froze at similar levels (**Fig. 4F**), indicating that 5-HT_2A_ receptor contributes to psilocybin’s effect on extinction retention. By contrast, psilocybin remained effective relative to vehicle in mice that were co-administered with WAY100635 (*p* = 0.025; **Fig. 4I**). For fear renewal on day 11, like **Fig. 1G**, vehicle + psilocybin mice froze less than vehicle + vehicle mice (*p* = 0.002; **Fig. 4D**). No effect of psilocybin was observed in MDL100907-treated mice, which froze at approximately control levels (**Fig. 4G**). Similarly, no effect of psilocybin was observed in WAY100635-treated mice for fear renewal (**Fig. 4J**). Collectively, these results provide evidence that psilocybin’s long-term effects on enhanced extinction relies on 5-HT_2A_ receptors, with the suppression of fear renewal in unfamiliar context additionally dependent on 5-HT_1A_ receptors.

## DISCUSSION

There are two main findings for this study. First, psilocybin enhances fear extinction, including longer term effects in extinction retention and fear renewal in a novel context. Second, the effect of psilocybin on fear extinction is dependent on dose, timing of administration, and contributions from 5-HT_2A_ and 5-HT_1A_ receptors.

We found a dynamic dose-response relationship for psilocybin-enhanced extinction, as 0.5, 1 and 2 mg/kg psilocybin all reduced conditioned freezing for extinction acutely, but only the 1 mg/kg dose showed persisting effects in extinction retention and fear renewal. The dose dependence may arise because fear extinction is a complex behavior moderated by concerted changes in attention, arousal, and memory, which often exhibit U-shaped dose-effect relationships^48,49^. Interestingly, a previous study assessing psilocybin’s impact on acoustic startle in rats came to a similar conclusion, identifying 1 mg/kg as the dose causing the greatest extent of behavioral change, while doses beyond 4 mg/kg failed to elicit lasting effects^50^. We explored potential sex difference in psilocybin’s dose-dependent behavioral effects. We detected statistical differences but did not proceed to post hoc analyses because of few animals per group when the data were fully segmented by sex and dose. Qualitatively, female mice had a narrower dose-response relationship with 1 mg/kg as the optimal dose, whereas male mice exhibited sensitivity to a wider range of psilocybin doses. These results highlight the importance of testing different doses and sexes when evaluating psilocybin’s effects using preclinical behavioral assays.

Psilocybin was only effective when administered prior to extinction, as injection before fear conditioning or following extinction testing yielded no behavioral change. The timing of administration, and by extension the context following drug administration, is therefore crucial for psilocybin. The timing requirement is consistent with previous studies investigating the effects of MDMA and low dose of scopolamine on fear extinction^15,51^, but differs sharply with ketamine which can accelerate fear extinction and suppress fear renewal even when administered well in advance^52^. Moreover, the behavioral domain may matter because prior work showed that psilocybin rescued stress-induced deficits in sucrose preference^53^ and learned helplessness^34^ when administered 24 hours prior to testing. This raises an intriguing question of whether certain psychedelics and tasks may require concurrent behavioral experience to produce their translationally important effects^54,55^. Thus, it is worth thinking beyond the simple picture of drug-evoked synaptogenesis and ask how these novel synapses are integrated into neural circuits in a behaviorally relevant manner for the appropriate context.

One caveat of the study is that although WAY100635 is a potent 5-HT_1A_ receptor antagonist^47^, it has the off-target effect of activating D_4_ receptors^56^. The results in the WAY100635 experiment may therefore be due in part to agonism of D_4_ receptors. That said, there are previous studies which corroborate the potential role of 5-HT_1A_ receptors in mediating psilocybin’s action. Mice lacking the 5-HT_1A_ receptor show heightened fear responses when encountering previously conditioned cues in novel contexts^57,58^. Attenuating 5-HT_1A_ receptor activity in novel contexts, but not familiar ones, blocks induction of long-term potentiation of medial perforant path inputs to the dentate gyrus^59^. Yohimbine has been extensively tested as a potential agent for enhancing fear extinction, and a potential target for its mechanism of action is 5-HT_1A_ receptors^60^. Moreover, our results agrees generally with prior studies showing the importance of 5-HT_2A_ receptors in mediating psychedelics’ effects on various fear-based assays^16,17,19,21,33,61^, but here more specifically we highlight the importance for extinction at the long time scale. Another caveat is that we rely on conditioned freezing as the sole readout of learned fear. This is advantageous because the measurement was automated to eliminate experimenter bias. However, the readout ignores other potential response strategies, which may be differentially employed by male and female rodents^36^. In the future, it will be useful to compare the effects of psilocybin on fear extinction retention using other readout methods and in other species, such as humans where MDMA has been tested in preliminary studies^62,63^.

In summary, this study shows that psilocybin enhances fear extinction and suppresses fear renewal long after the drug has been administered. Dose, context, and serotonin receptors are contributing factors to psilocybin’s effects. The suppression of fear renewal is particularly relevant for translation because this is an unmet need in the clinic. Thus, the findings provide a detailed characterization of psilocybin’s effects on behavioral processes relevant to prolonged exposure therapy. Moreover, the work adds further preclinical support for investigating the utility of psilocybin as a pharmacological adjunct for extinction-based therapy for PTSD.

## Acknowledgements

We thank Renu Sah and Michael Fanselow for advice on the behavioral studies. Psilocybin was provided by Usona Institute’s Investigational Drug & Material Supply Program; the Usona Institute IDMSP is supported by Alexander Sherwood, Robert Kargbo, and Kristi Kaylo in Madison, WI. This work was supported by NIH grants R01MH128217 (A.C.K.); One Mind – COMPASS Rising Star Award (A.C.K.); Rawlings Cornell Presidential Research Scholars program (C.M.L.).

## Contributions

S.C.W. and A.C.K planned the study. S.C.W. performed experiments. C.M.L. and A.M.K. assisted with the experiments. S.C.W. analyzed the data. S.C.W. and A.C.K. drafted the manuscript. All authors reviewed the manuscript before submission.

## Competing interests

A.C.K. has served as a scientific advisor for Empyrean Neuroscience, Freedom Biosciences, and Psylo. A.C.K. has received research support from Intra-Cellular Therapies. The other authors report no financial relationships with commercial interests.

